# Scopoletin Augments DJ1/Nrf2 Signalling and Prevents Protein Aggregation in Parkinson’s disease

**DOI:** 10.1101/260521

**Authors:** S Narasimhan Kishore Kumar, Jayakumar Deepthy, Velusamy Prema, Srinivasan Ashokkumar, Mohan Thangarajeswari, Ravi Divya Bhavani, Uthamaraman Saraswathi, Sathyamoorthy Yogesh Kanna, Namakal S. Rajasekaran, Periandavan Kalaiselvi

## Abstract

Given the role of oxidative stress in PD pathogenesis and off-target side effects of currently available drugs, several natural phytochemicals seems to be promising in the management of PD. Here, we tested the hypothesis that scopoletin, an active principle obtained from *Morinda citrifolia* (MC), efficiently quenches oxidative stress through DJ-1/Nrf2 signalling and ameliorates rotenone-induced PD. Despite reducing the oxidative stress, administration of MC extract (MCE) has lessened the protein aggregation as evident from decreased levels of nitrotyrosine and α-Synuclein. *In vitro* studies revealed that scopoletin lessened rotenone-induced apoptosis in SH-SY5Y cells through preventing oxidative injury. Particularly, scopoletin markedly up-regulated DJ-1, which then promoted the nuclear translocation of Nrf2 and transactivation of antioxidant genes. Furthermore, we found that scopoletin prevent the nuclear exportation of Nrf2 by reducing the levels of Keap1 and thereby enhancing the neuronal defence system. Overall, our findings suggest that scopoletin acts through DJ-1 mediated Nrf2 signalling to protect the brain from rotenone-induced oxidative stress and PD. Thus, we postulate that scopoletin could be a potential drug to treat PD.

## 1. Introduction

Although the causal mechanisms of Parkinson disease (PD) remain elusive, excess production of reactive oxygen species (ROS), mitochondrial dysfunction, neuro-inflammation and environmental toxins are reported to promote the loss of dopaminergic neurons in PD [1]. Oxidative stress has been shown to induce misfolding, aggregation and accumulation of such aggregates leading to the pathogenesis of many neurodegenerative diseases including PD [2]. Intracellular inclusions known as Lewy Bodies (LBs) are regarded as a hallmark of common pathological manifestation in both familial and sporadic PD patients with α-Synuclein (α-Syn) serving as the main component of LB [3]. α-Syn is natively unfolded and is prone to form fibrils during oxidative stress [4], indicating that redox signalling may play a significant role in the aggregation of α-Syn.

Previous studies have reported that loss of antioxidant defence aggravates PD progression [5, 6]. A key example includes DJ-1/PARK7, a molecular chaperone known to regulate Keap1-Nrf2 signalling, which is the primary sensor for reactive electrophiles activating Nrf2 nuclear translocation and transactivation of the antioxidant response element (ARE) in a battery of cytoprotective genes facilitating protection from oxidative stress pathogenesis [7] including experimental models of PD [8]. Thus, pharmacological activation of the Nrf2 in the brain is likely to preserve neuronal health. Therefore, exploration for therapeutic compounds with lesser neurotoxic effects that activates Nrf2 signaling would be promising to treat PD.

In this context, identifying potential principles from medicinal plants would be ideal as plant extracts have been reported to have several therapeutic benefits, due to the synergistic effect of various natural ingredients [9]. However, such plant sources have not been in clinical practice or in global market due to the lack of scientific validation. Here, we hypothesized that *Morinda citrifolia* extract (MCE) and scopoletin prevents rotenone-induced oxidative stress and apoptosis through activation of DJ1/Nrf2 signalling, thereby ameliorates PD in rats.

## 2. Materials and Methods

### 2.1. Animals, Intra-Nigral rotenone infusion and treatment

Adult male Sprague-Dawley rats were used in this study. All experiments were performed in accordance with the guidelines approved by the Institutional Animal Ethical Committee *(IAEC No. 01/09/12).* Rats were divided into five groups (n=10/gp.). Group I served as control, while Group II to V were subjected to stereotaxic surgery. Group II served as sham controls and groups III-V were stereotaxically infused with rotenone to induce Parkinsonism. Briefly, rats were anaesthetized with ketamine hydrochloride and xylazine (80 mg/kg and 10mg/kg; i.p.) and placed on a small animal stereotaxic frame (Stoelting, IL, USA). Rotenone dissolved in DMSO (1μ§/1μΓ) was infused into the right Ventral Tegmental Area (VTA, Anterior-Posterior (AP): 5.0 mm; Laterally (L): 1.0 mm; Dorso-Ventral (DV): 7.8 mm) and into the right Substantia Nigra Pars compacta (SNPc, AP: 5.0 mm; L: 2.0 mm; DV: 8.0 mm) at a flow rate of 0.2 μl/min using a Hamilton 26 gauge needle [10, 11]. The infusion needle was left in place for additional five minutes for complete diffusion of the drug. Sham controls were infused with DMSO and polyethylene glycol in the ratio of 1:1 during stereotaxic surgery. Two weeks post-surgery, rats in Group IV and V were treated with Levodopa (LD, 100mg/kg with 25mg/kg benserazide [12]) and ethyl acetate extract of *Morinda citrifolia* fruit (MCE, 150mg/kg body weight), respectively, for the next 30 days. To determine the efficiency of intranigral-infusion of rotenone, animals were subjected to behavioural analysis [11].

### 2.2. Statistical Analysis

Data are presented as mean ± standard error of mean (SEM) of the results obtained from the average of at least three to six independent experiments. Results were analysed by one-way analysis of variance (ANOVA) using the SPSS software package for Windows (Version 20.0; SPSS Inc., Chicago, IL, USA) and p values were determined using the Student-Newman-Keuls and least significant difference post hoc test. Differences among means were considered statistically significant when the p value was less than 0.05.

## 3. Results

### 3.1 Impact of MCE on nigrostriatal Tyrosine Hydroxylase (TH) immunoreactivity

Immunohistochemical localization of TH positive neurons in the striatum as well as the sunstantia nigra pars compacts (SNpc) of control and experimental rats are presented in **Fig 1A**. Rotenone infused SNpc showed less number of TH positive cells (43% reduction) as compared to control animals. MCE treatment in these animals restored the TH+ cells. **(Fig. 1B)**.

**Figure 1:**
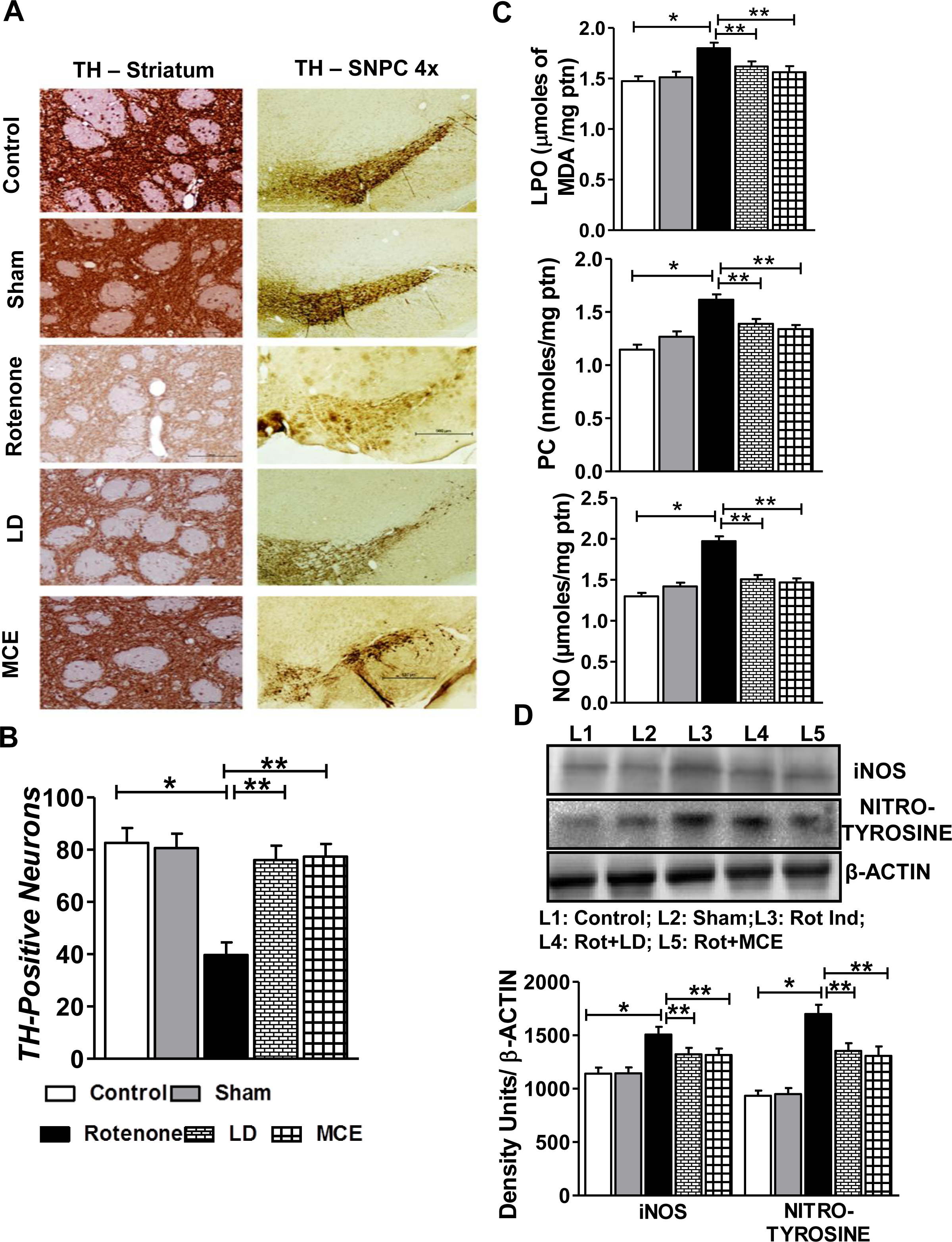
**(A) Photographic representation of the TH-positive neurons in the striatum and in SNPc. (B) Quantification of relative intensity of TH staining** in striatum using the Densitometry protocol through ImageJ was performed to substantiate the potentials of salvaging activity of MCE. **(C) Impact of MCE on rotenone induced oxidative stress:** Values are expressed as mean ± SEM for six animals in each group. **(D) Immunoblot analysis of iNOS and nitrotyrosine and representative densitometry quantification**. Statistical significance (p< 0.05) was calculated by Student-Newman-Keuls and least significant difference post hoc test, where *represents Control Vs other groups, **represents Rotenone vs LD, MCE.

### 3.2. MCE prevents rotenone induced oxidative stress by augmenting the antioxidant defence systems in experimental PD rats

Analyses of various oxidative stress markers indicated that MCE was an efficacious treatment to reduce oxidative stress in rotenone-induced PD rats. The levels of nitric oxide (NO), lipid peroxidation (LPO) and protein carbonyls (PC) were significantly increased (p<0.05) in the striatum of rotenone-infused PD rats and this was blunted in response to MCE treatment (**Fig. 1C**). A significant decline (p < 0.05) in the activities of antioxidant enzymes, namely, superoxide dismutase (SOD), catalase, glutathione peroxidase (Gpx), glutathione reductase (GR) and glutathione-S-Transferase (GST) along with declined GSH was observed **(Table 1)** in the rotenone induced PD rats. However, upon supplementing with MCE, a significant augmentation in the activities of SOD, CAT and GST were observed with a maximum improvement in the SOD activity (36%) and reversed the GSH levels. As we observed a significant decline in Nrf2 protein levels, we further assayed transcript and protein levels for Nrf2 gene targets. Consistent with our observation of oxidative insult and Nrf2 pathway antagonism, rotenone treatment was associated with a prominent decline in the mRNA and protein **(Fig. S1A-B)** levels of several ARE targets with a maximum decrease being observed in HO-1 (45%). MCE treatment significantly enhanced the levels of these proteins both at the transcriptional and translational levels by an average of 40%. The improved protein levels are reflected in the activities of these enzymes **(Table 1)**.

**Table 1:** Impact of MCE on antioxidant defense system and the activities of Nrf2/ARE downstream enzymes. Values are expressed as mean ± SEM for six animals in each group. Units: SOD - units/mg protein. One unit is equal to the amount of enzyme that inhibits the pyrogallol auto-oxidation by 50%; CAT - μmoles of H_2_O_2_ consumed/min/mg protein; GPx - μg of GSH consumed /min/mg protein; GR - nmoles of NADPH oxidized/min/mg protein; GST - μ moles of CDNB conjugate formed/minute/mg protein; GSH - nmoles/mg protein; γGCLC - millimoles of NADH oxidized/min/mg protein; NQO1 - nmoles DCPIP utilized/min/mg protein; HO-1 - nmoles of bilirubin/hr/mg protein. Statistical significance (p< 0.05) was calculated by Student-Newman-Keuls and least significant difference post hoc test, where *represents Control Vs other groups, **represents Rotenone vs LD, MCE.

| **Parameter** | **Control** | **Sham**<br>**Control** | **Rotenone**<br>**Induced** | **LD** | **MCE** |
| --- | --- | --- | --- | --- | --- |
| **SOD** | 0.58 ± 0.018 | 0.52 ± 0.017 | 0.33 ± 0.015* | 0.41 ± 0.016** | 0.45 ± 0.014** |
| **CAT** | 17.43 ± 0.60 | 16.36 ± 0.36 | 12.13 ± 0.38* | 14.21 ± 0.32** | 15.22 ± 0.41** |
| **GPx** | 8.25 ± 0.37 | 8.22 ± 0.32 | 5.66 ± 0.23* | 6.59 ± 0.31** | 6.64 ± 0.20** |
| **GR** | 5.16 ± 0.25 | 5.12 ± 0.22 | 3.90 ± 0.15* | 4.47 ± 0.19** | 4.48 ± 0.15** |
| **GST** | 9.08 ± 0.394 | 9.01 ± 0.468 | 6.38 ± 0.271* | 7.75 ± 0.298** | 7.86 ± 0.241** |
| **GSH** | 1.93 ± 0.055 | 1.76 ± 0.048 | 1.28 ± 0.058* | 1.61 ± 0.059** | 1.65 ± 0.062** |
| **γGCL** | 2.54 ± 0.084 | 2.46 ± 0.083 | 1.35 ± 0.051* | 1.90 ± 0.071** | 2.01 ± 0.068** |
| **NQO1** | 5.88 ± 0.193 | 5.85 ± 0.193 | 3.15 ± 0.111* | 4.57 ± 0.151** | 4.82 ± 0.153** |
| **HO-1** | 0.81 ± 0.026 | 0.80 ± 0.026 | 0.48 ± 0.016* | 0.63 ± 0.021** | 0.65 ± 0.018** |

### 3.3. MCE protects from rotenone-induced protein aggregation

Rotenone infused Parkinsonian rats showed a significant (p<0.05) increase in the protein levels of iNOS and nitrotyrosine when compared with controls **(Fig. 1D)**. Treatment with MCE significantly ameliorated rotenone-mediated NO production through diminishing the levels of iNOS. As a result, we also found that MCE was particularly efficient in blocking the formation of nitrotyrosine adducts, which are the primary events in the process of protein aggregation that occurs in response to oxidative insults [2]. To further confirm the suppression of protein aggregation by MCE, we analysed the aggregation of α-Synuclein (α-Syn) by immuno-staining (**Fig. 2A-B**). Rotenone infused Parkinsonian rats exhibited a significant (p<0.05) increment in the protein levels of α-Syn by 79% **(Fig. 2C)**, which further aggravated the aggregation of α-Syn in these rats. While immunostaining showed a significant increase in α-Syn aggregation in the striatum of rotenone-induced rats which correlates with the increase in the nitrotyrosine levels, MCE treatment abolished these changes and significantly reduced the aggregation of α-Syn **(Fig. 2A-B)**, suggesting suppression of oxidative stress is likely through MCE mediated augmentation of antioxidative signaling.

**Figure 2:**
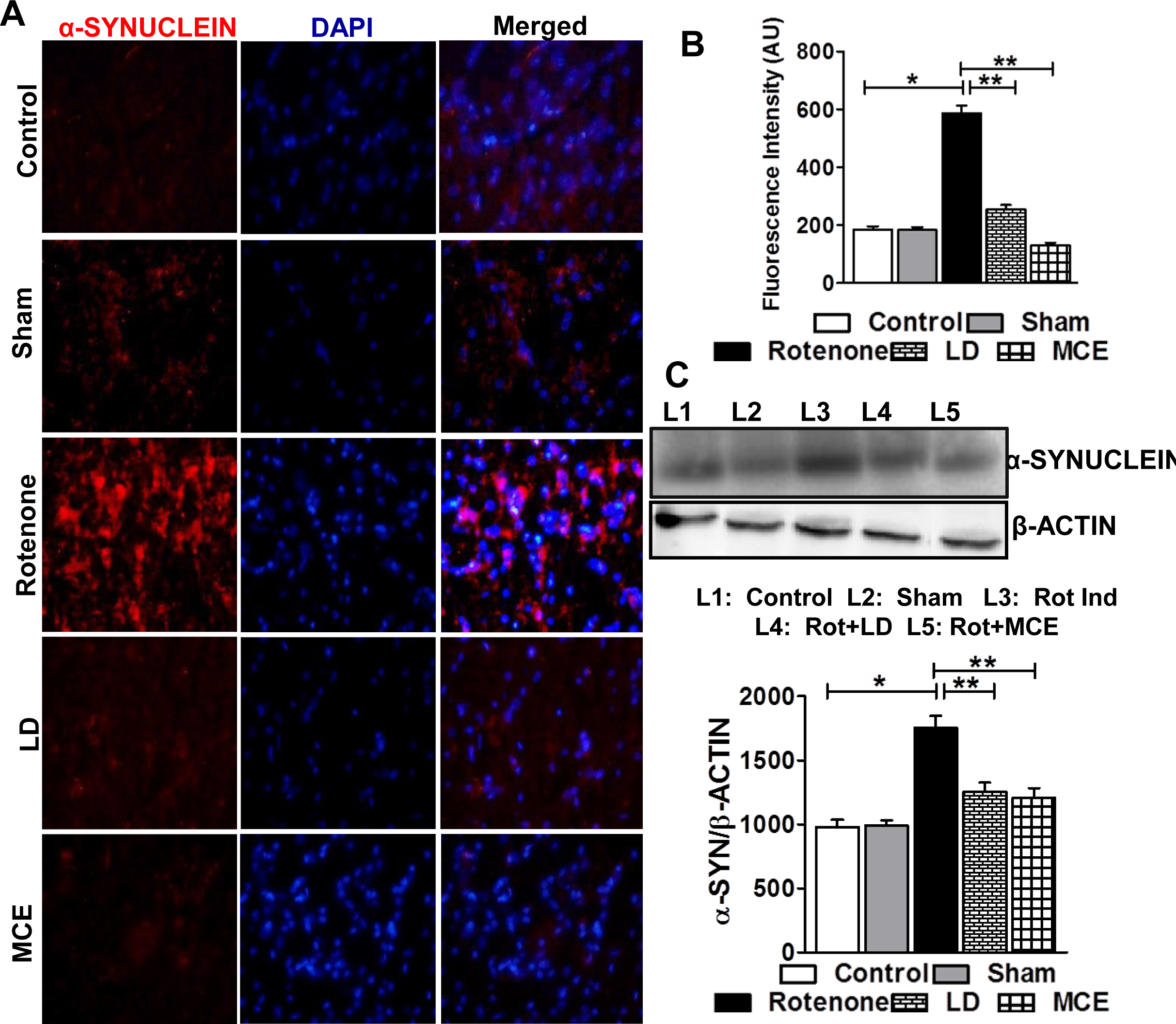
Rotenone induced aggregation of α-Synuclein (α-Syn) measured by immunofluorescence analysis. **(A)** The cells were visualized using fluorescence microscopy and images captured using 20x magnification. The control and sham group show very low level of expression, however the expression is punctate and high in the rotenone induced group, the expression is meagre in levodopa, on the contrary expression in MCE groups is comparable with that of the control group. **(B)** Relative fluorescence intensity was calculated. **(C)** Immunoblot analysis of α-Syn and representative densitometry quantification. Statistical significance (p< 0.05) was calculated by Student-Newman-Keuls and least significant difference post hoc test, where *represents Control Vs other groups, **represents Rotenone vs LD, MCE.

### 3.4. MCE induces Nrf2/ARE pathway and supress rotenone induced oxidative stress

Downregulation of the Nrf2/ARE pathway exacerbates oxidative stress which potentiates dopaminergic degeneration and pathogenesis of PD [13]. Thus, we further analysed the levels of Nrf2 and its interacting proteins to test the hypothesis that MCE-mediated augmentation of antioxidative system occurs through activation of Nrf2/ARE signalling. Immunoblotting analysis revealed that MCE-treatment significantly rescued the levels of Nrf2 and reversed the rotenone-induced increase in Keap1 and Cullin3 (**Fig. 3A)**. Next, we extended our immunoblotting analyses of total protein expression to nuclear and cytosolic fractions (**Fig. 3B)**. Our results confirmed that rotenone infusion significantly repressed the activation of Nrf2 as nuclear Nrf2 protein decreased in rotenone versus control rats. Hence, rotenone-infusion not only decreases total Nrf2 protein levels, but also impairs its nuclear translocation by augmenting cytosolic Keap1 levels. As such, supplementation with MCE significantly augmented the nuclear translocation of Nrf2 as evident from increased levels of nuclear Nrf2 (83%) by decreasing the nuclear Keap1 levels when compared with rotenone alone infused rats.

**Figure 3:**
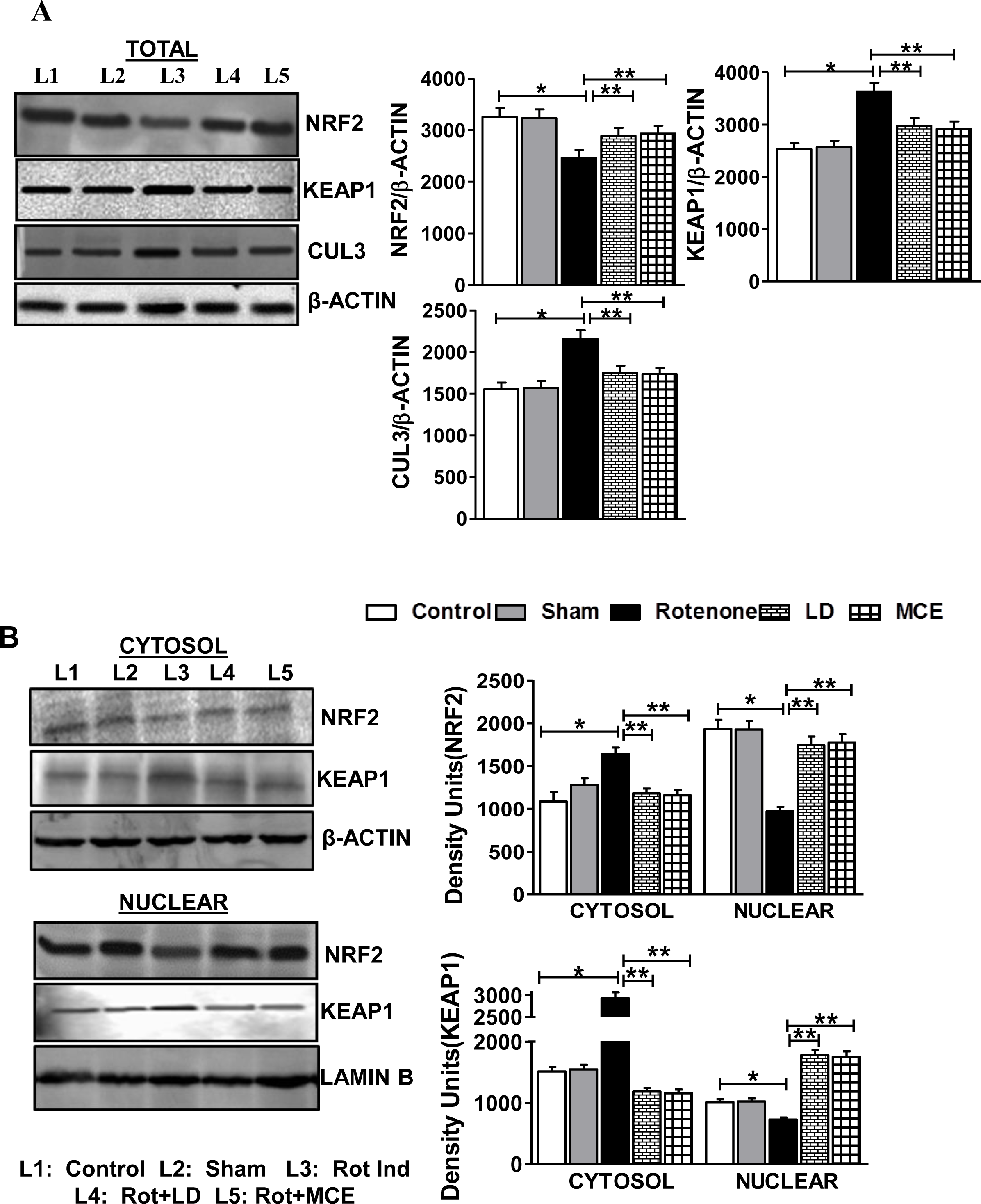
Immunoblot Analysis of Nrf2 and its negative regulators Keap1 and cullin3 in the striatum of rotenone induced PD rats. **(A)** The total protein levels of Nrf2, Keap1 and Cullin3. **(B)** The protein levels of Nrf2 and Keap1 in cytosolic and nuclear compartment of striatal neuronal cells. Statistical significance (p< 0.05) was calculated by Student-Newman-Keuls and least significant difference post hoc test, where *represents Control Vs other groups, **represents Rotenone vs LD, MCE.

### 3.5. Scopoletin-mediated neuroprotective effects were associated with the translocation of nuclear p40Nrf2 and upregulation of DJ-1

SH-SY5Y cells undergoing various stages of apoptosis (early, mid-stage and late stage) were analyzed by flow cytometry using Annexin V and propidium iodide (PI) dual staining. Treatment of SH-SY5Y cells with rotenone (500nM for 24 h [14]) resulted in 40% cell-death, which was attenuated by pre-treating the cells with 30μΜ scopoletin for 3 hrs **(Fig. 4A-B)**. These data indicate that scopoletin protects SH-SY5Y from rotenone-induced cell death. As scopoletin prevented rotenone-induced cell death, we further assessed the regulation of Nrf2 signalling **(Fig. 4C)**. Along these lines, SH-SY5Y cells were pre-treated with 30μΜ scopoletin for 3 hours and both the un-phosphorylated (total/cytosolic form) and the serine 40 phosphorylated Nrf2 (pNrf2S40) levels were measured by immunoblotting. Consistent with animal studies demonstrating that MCE improves nuclear levels of Nrf2, scopoletin augmented the nuclear translocation of Nrf2 as evident from the increased nuclear levels of pNrf2 (S40) **(Fig. 4D)**. This increase in the nuclear levels of Nrf2 may be attributed to the phosphorylation of Nrf2 by PKC-δ which is also augmented upon pre-treating with scopoletin. Concomitant with the animal studies, both the total and nuclear levels of Keapl were significantly elevated in rotenone alone treated SH-SY5Y cells when compared with untreated control cells. Furthermore, rotenone treatment also increased the levels of E3 ubiquitin ligase Cullin3 by 46%.

**Figure 4:**
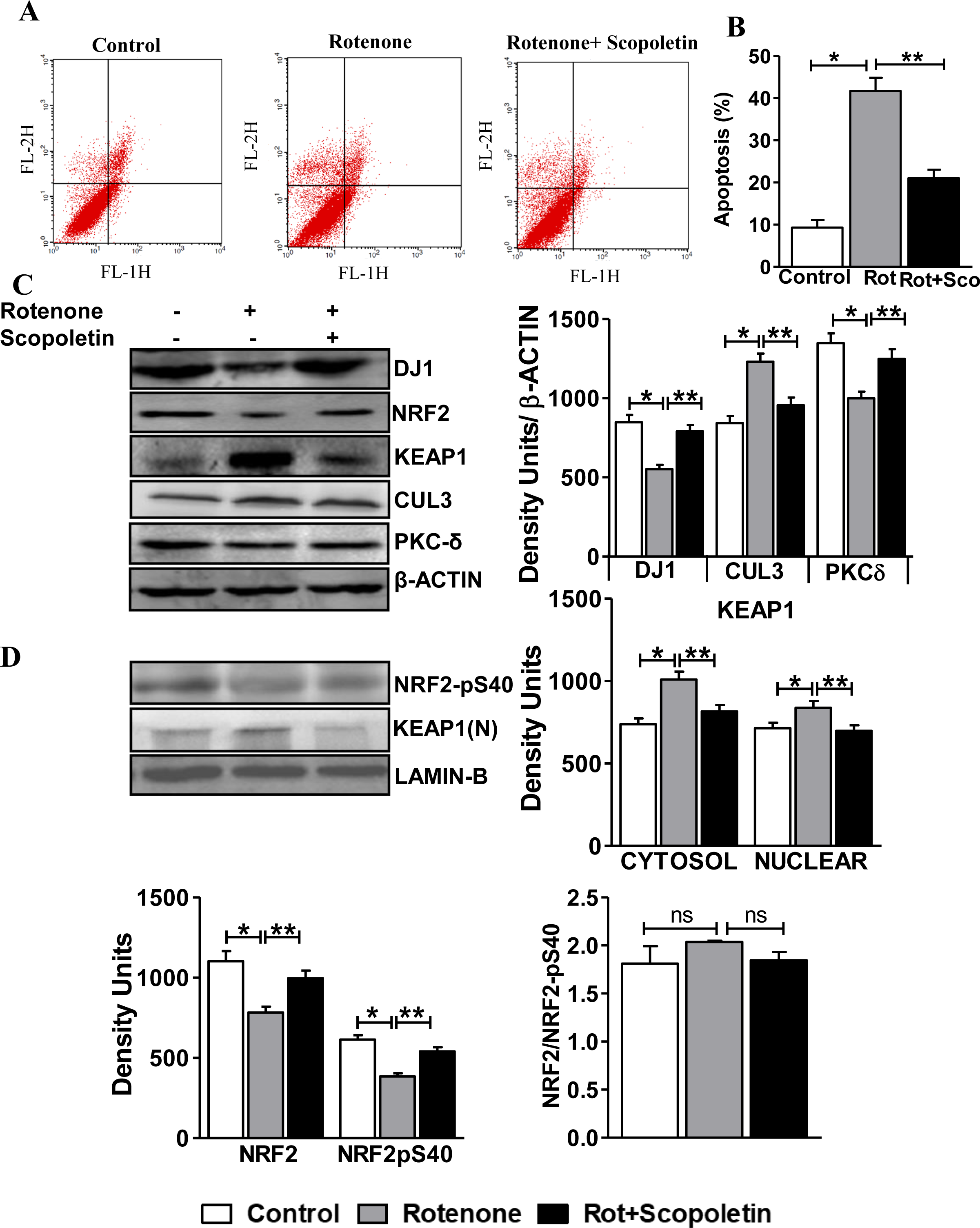
(A) Rotenone induced apoptosis is counteracted by scopoletin in SH-SY5Y cells. At the end of experiment, cells were stained with annexin-V/FITC and read immediately by flow cytometry to measure the extent of apoptosis; X-axis FL-1H denotes the FITC and Y-axis FL-2H denotes the PI**. (B)** Quantification of percentage dead cells. **(C)** Immunoblot analysis of several proteins involved in Nrf2/ARE pathway is performed in SH-SY5Y cells treated with/without rotenone and scopoletin **(D)** Immunoblot analysis of phospho (S40) Nrf2 and nuclear Keap1 in SH-SY5Y cells. Statistical significance (p< 0.05) was calculated by Student-Newman-Keuls and least significant difference post hoc test, where *represents Control Vs other groups, **represents Rotenone vs Scopoletin.

Recent reports indicate that nuclear translocation of Nrf2 is not only mediated by phosphorylation by PKC-δ but also by intact DJ1, which binds and stabilizes the Nrf2 and favours its translocation to the nucleus [15]. While rotenone decreases the levels of DJ-1, scopoletin pre-treated cells showed an augmented expression of DJ-1 (46%), suggesting that DJ-1 may be vital for scopoletin-mediated neuroprotective effects. From these observations, it is clear that scopoletin augments the Nrf2/ARE pathway by increasing levels of DJ-1 and concomitantly prevents the cytosolic degradation of Nrf2.

## 4.0. Discussion

Parkinson's disease (PD) is the second most common progressive neurodegenerative disorder with increased oxidative stress being the key player of the pathogenesis. Therefore, increasing interest has been focused on identifying dietary supplements and phytoconstituents that can ameliorate oxidative stress mediated protein aggregation and cell death, thereby reversing the multi-faceted pathophysiological events underlying PD. Here, we investigated whether *Morinda citrifolia* (noni) fruit extract (MCE) attenuates rotenone-induced oxidative stress by activating the Nrf2-dependent antioxidant response. We elucidated the mechanism of action for scopoletin, a major compound present in the MCE, *in vitro* using SH-SY5Y cells and identified that the therapeutic effect of scopoletin is facilitated through activation of the DJ-1/Nrf2/ARE signalling cascade.

Recent reports have demonstrated that an imbalance of nitric oxide (NO) signalling underlies the incidence of oxidative stress in PD [2, 16]. Here, we have shown that MCE supplementation significantly abrogates rotenone mediated increase in NO and iNOS in the striatum of PD induced rats. These results provide evidence for the antioxidative potential of MCE in the brain. Although levodopa (LD), used as a positive control to treat rotenone-infused mice, seems to be protective in the current study, its chronic use might be detrimental as previous studies have shown that nitric oxide levels were increased in LD treated PD patients [17]. Therefore, use of a natural antioxidant like *Morinda citrifolia* or coadministered with LD might hold promising outcomes in PD.

As protein aggregation is a common event underlying neurodegenerative diseases including PD, wherein α-Synuclein comprises the bulk of Lewy bodies [18, 19], we assessed whether the antioxidant capacity of MCE could supresses α-Synuclein aggregation. Indeed, MCE treatment attenuated the increase in α-Synuclein expression and aggregation following rotenone exposure. Consistent with the relationship between NO and α-Synuclein aggregate formation, the cytoprotective effects of MCE may be mediated though its impact on NO availability and tyrosine nitration [20].

To combat oxidative insult, the intracellular defense is maintained through a variety of antioxidant enzymes and non-protein thiols such as glutathione (GSH) [21, 22]. In the current study, rotenone infusion resulted in a significant decline in striatal GSH content, and diminished the enzymatic activities of several key antioxidants. Notably, MCE supplementation restored antioxidant capacity and rescued GSH levels in the striatum of rotenone-infused PD rats. These findings indicate that MCE exhibits strong antioxidative potential that effectively combats oxidative stress. Because Nrf2 serves as the master transcriptional activator of the antioxidant response element (ARE), we wondered whether MCE mediated antioxidant defense occurred through enhanced Nrf2 signalling. Although rats stereotaxically infused with rotenone exhibited a significant decline in Nrf2-dependent ARE induction, MCE treatment stabilized Nrf2 activity resulting in an increased expression of downstream targets γGCLC, NQO1 and HO-1.

After demonstrating efficacy for MCE, we proceeded to investigate the therapeutic potential of its active constituent scopoletin *in vitro.* Scopoletin pre-treatment effectively attenuated rotenone mediated apoptosis in SH-SY5Y cells. In light of our observations on MCE mediated Nrf2 activation, we examined the capacity for scopoletin to influence the Nrf2 signalling. In the present investigation, the total Nrf2 protein was significantly reduced in rotenone induced rats and SH-SY5Y cells while the Keap1 and cullin3, negative regulators of Nrf2, were markedly elevated. Either an increase in Keap1 or high levels of oxidative stress may promote the degradation of Nrf2, although the molecular mechanism for the latter is uncertain. Our results indicate that MCE and scopoletin administration suppresses Keap1 and Cullin3, thereby preventing degradation of Nrf2 *in vivo* and *in vitro*, respectively. Our profiling of ARE-dependent transcript and protein expression as well as enzymatic activities implicated Nrf2 activation in mediating the cytoprotective effects of MCE. Indeed, our analysis of Nrf2 nuclear translocation and phosphorylation status confirmed this hypothesis as MCE increased nuclear Nrf2 protein in the striatum of rotenone infused PD rats. Similarly, scopoletin is capable of supporting Nrf2 activation, as evident from the increased levels of phospho-Nrf2, and PKC-δ that phosphorylates serine 40 of Nrf2 in SH-SY5Y cells exposed to rotenone.

Previous reports have shown that DJ-1 stabilizes Nrf2 and promotes its nuclear translocation [25]. Intriguingly, mutations in DJ-1 are associated with risk of developing PD [26]. Hence, we further evaluated the effect of scopoletin on DJ-1 levels in SH-SY5Y cells. We found that pre-treating SH-SY5Y cells with scopoletin rescued DJ-1 protein levels, subsequently conferring protection against rotenone induced oxidative stress. Therefore, Nrf2 stabilization and activation underlies the therapeutic potential for MCE and scopoletin to combat oxidative stress during PD pathogenesis.

## 5. Conclusion

Our findings provide evidence that MCE prevented α-synuclein aggregation through augmenting Nrf2-antioxidant signalling and suppressing oxidative stress. Notably, scopoletin, the active component from MCE seems to be responsible for stabilizing Nrf2/ARE pathway by augmenting the phosphorylation of Nrf2 and its nuclear translocation, in a DJ-1-dependent manner. Thus, we propose that DJ-1 might be a potential target for scopoletin-based therapeutic strategy against neurodegenerative diseases.

## Abbreviations

DMSO: Dimethyl Sulfoxide
F-12K: Kaighn's Modification of Ham's F-12 Medium
FBS: Fetal bovine serum
GSH: Reduced Glutathione
HO-1: Heme Oxygenase-1
Keap1: Kelch-like erythroid cell derived protein with CNC (Cap “n” Collar) homology (ECH) protein1
LD: Levodopa
LPO: Lipid peroxidation
MCE: Ethyl acetate extract of *Morinda citrifolia* fruit
NO: Nitric oxide
NQO1: NAD(P)H: Quinone Oxidoreductase 1
Nrf2: Nuclear factor erythroid 2-related factor 2
PC: Protein Carbonyl
PD: Parkinson’s disease
PKC-δ: Protein kinase C delta
ROS: Reactive oxygen species
Rot Ind: Rotenone induced
SH-SY5Y: neuroblast-like subclone of SK-N-SH
SNpc: Substantia nigra pars compacta
VTA: Ventral Tegmental Area
γGCLC: Catalytic subunit of Gamma Glutamate Cysteine Ligase

## Acknowledgments

The financial assistance to Dr. S Narasimhan Kishore Kumar, Department of Medical Biochemistry from Indian Council of Medical Research in the form of ICMR Senior Research Fellow (SRF), New Delhi, Government of India is gratefully acknowledged.

Authors greatly appreciate Dr. Gobinath Shanmugam for assisting with art work (Figures) and Mr. Justin Quiles for his wonderful editorial assistance.

## Supplemental Figures

**Supplemental Figure 1: Increased Nrf2 regulated antioxidant genes in response to MCE supplementation in rotenone induced PD rats. (A)** The mRNA levels of Nrf2 downstream genes γGCLC, NQO1 and HO-1. **(B)** The protein levels of γGCLC, NQO1 and HO-1. Statistical significance (p< 0.05) was calculated by Student-Newman-Keuls and least significant difference post hoc test, where *represents Control Vs other groups, **represents Rotenone vs LD, MCE.

